## Supplementary Figures 1-9 for "Structural dynamics underlying gating and regulation in IP_3_R channel"

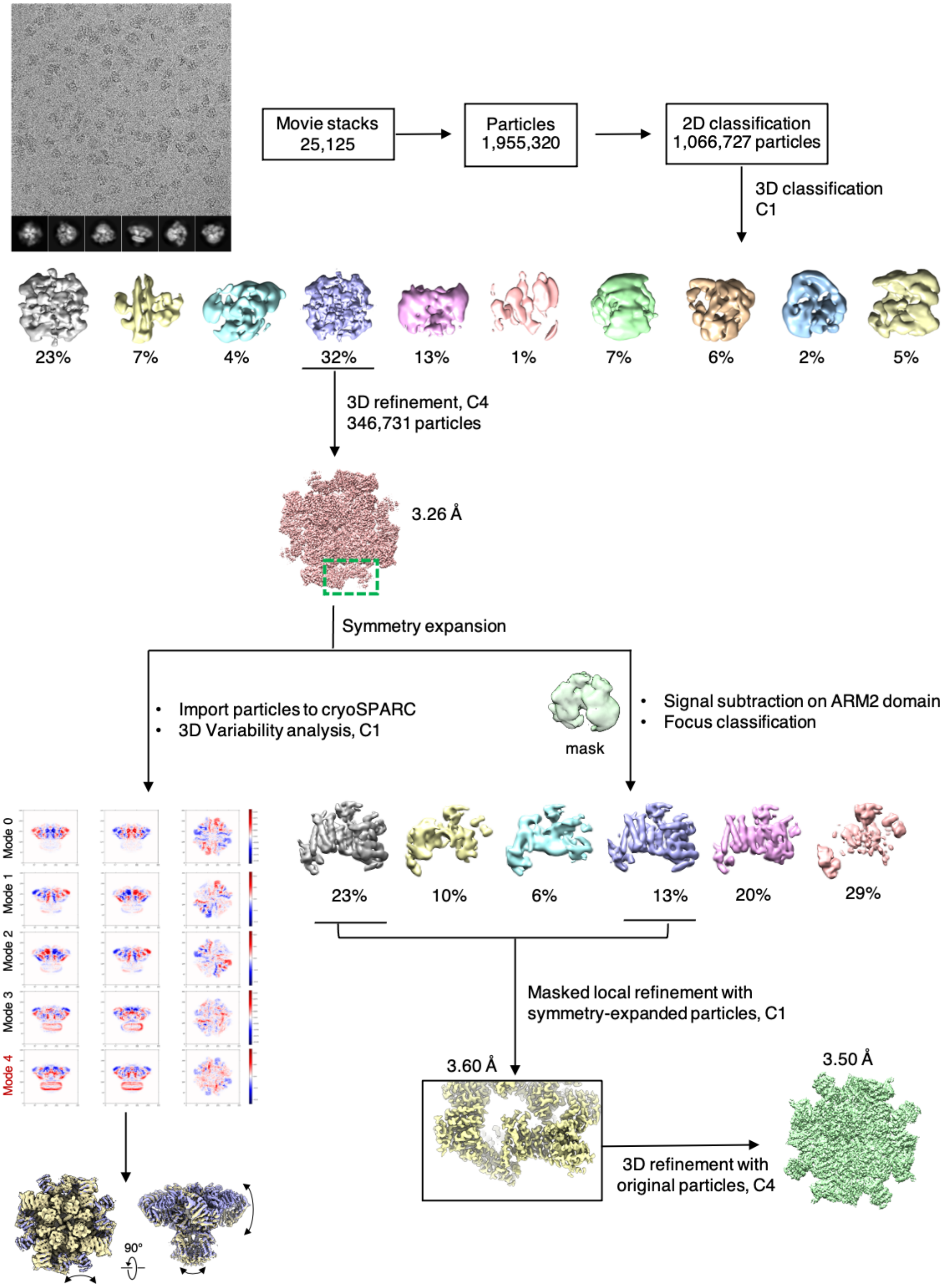


**Supplementary Figure 1. Cryo-EM processing workflow for Ca-IP$_3$R1.**

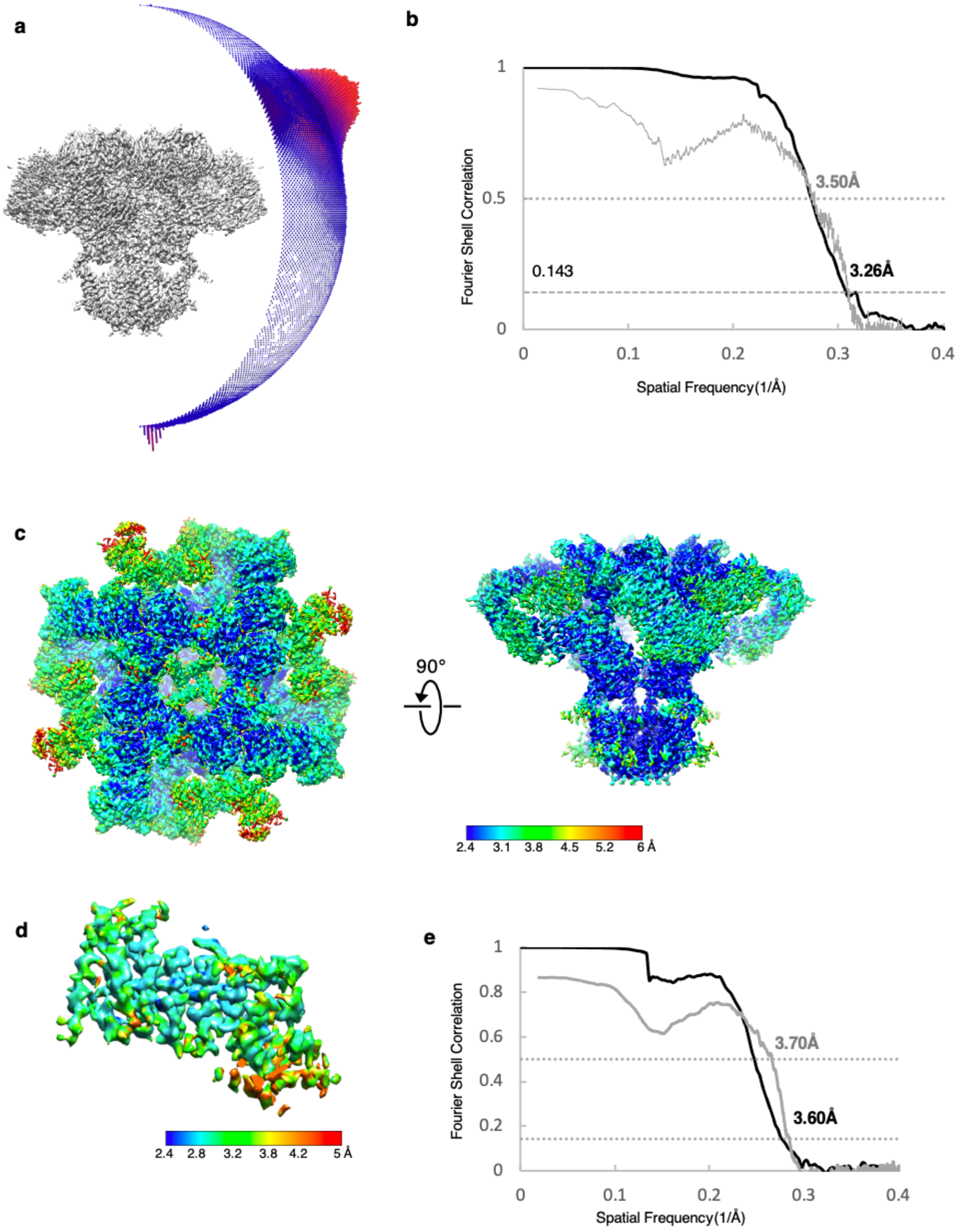


**Supplementary Figure 2. Characteristics of cryo-EM reconstructions for Ca-$IP_3R1$. a**, Euler angle distribution of particles contributing to the final reconstruction with larger red cylinders representing orientations comprising more particles. **b**, Resolution estimation of the cryo-EM density maps. Black – gold-standard Fourier shell correlation (FSC) curves, showing the overall nominal resolutions of 3.26 Å at 0.143 criterion; gray – Map-model FSC plot calculated by Phenix. **c**, Cryo-EM map colored based on local resolution. **d**, Focused 3D refined ARM2 map colored based on local resolution. **e**, Resolution estimation of the focused 3D refined ARM2 domain density map. Black – gold-standard Fourier shell correlation (FSC) curves, showing the overall nominal resolutions of 3.6 Å at 0.143 criterion; gray - Map-model FSC plot calculated by Phenix.

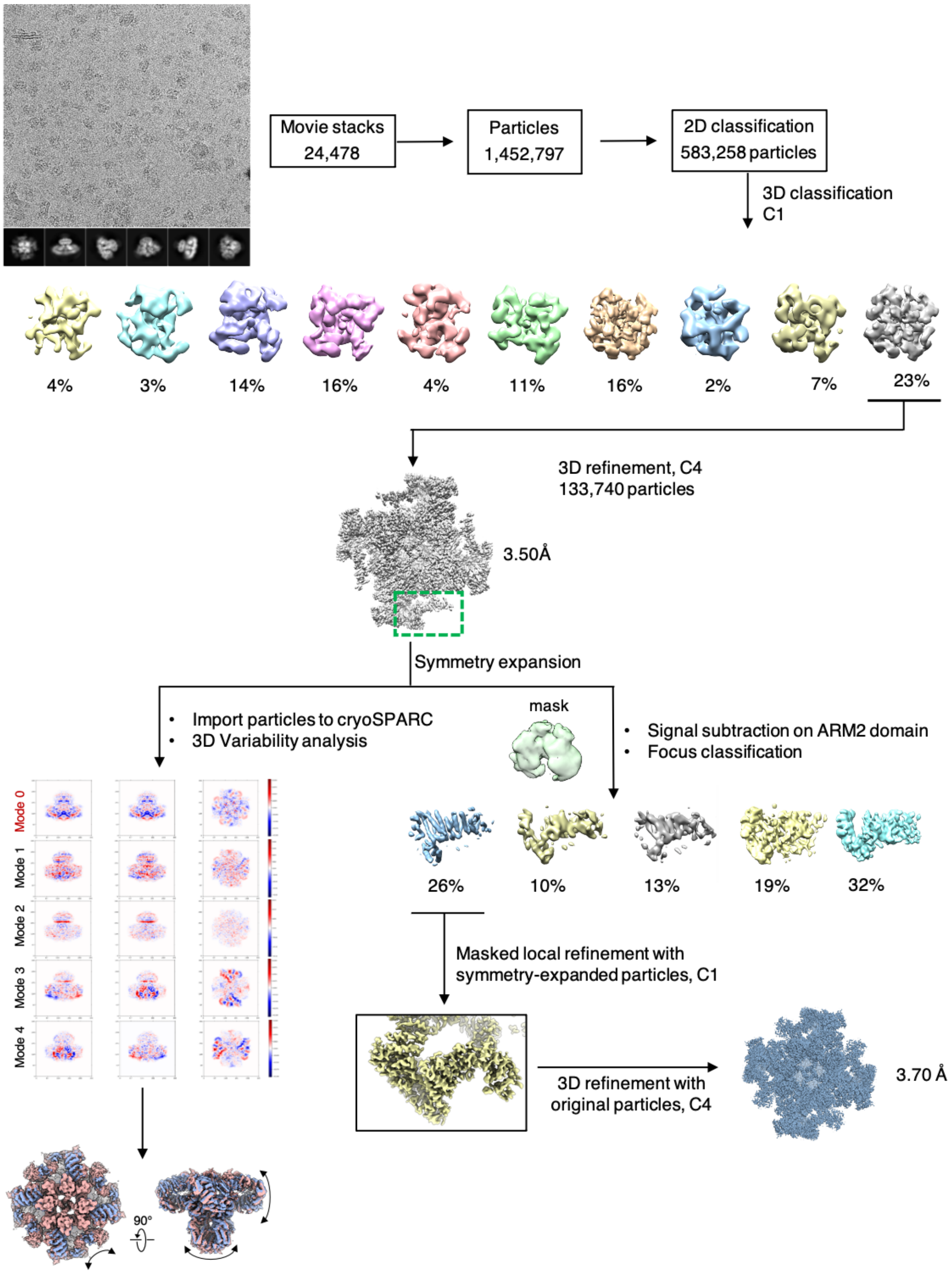


**Supplementary Figure 3. Cryo-EM processing workflow for CIA-$IP_3R1$.**

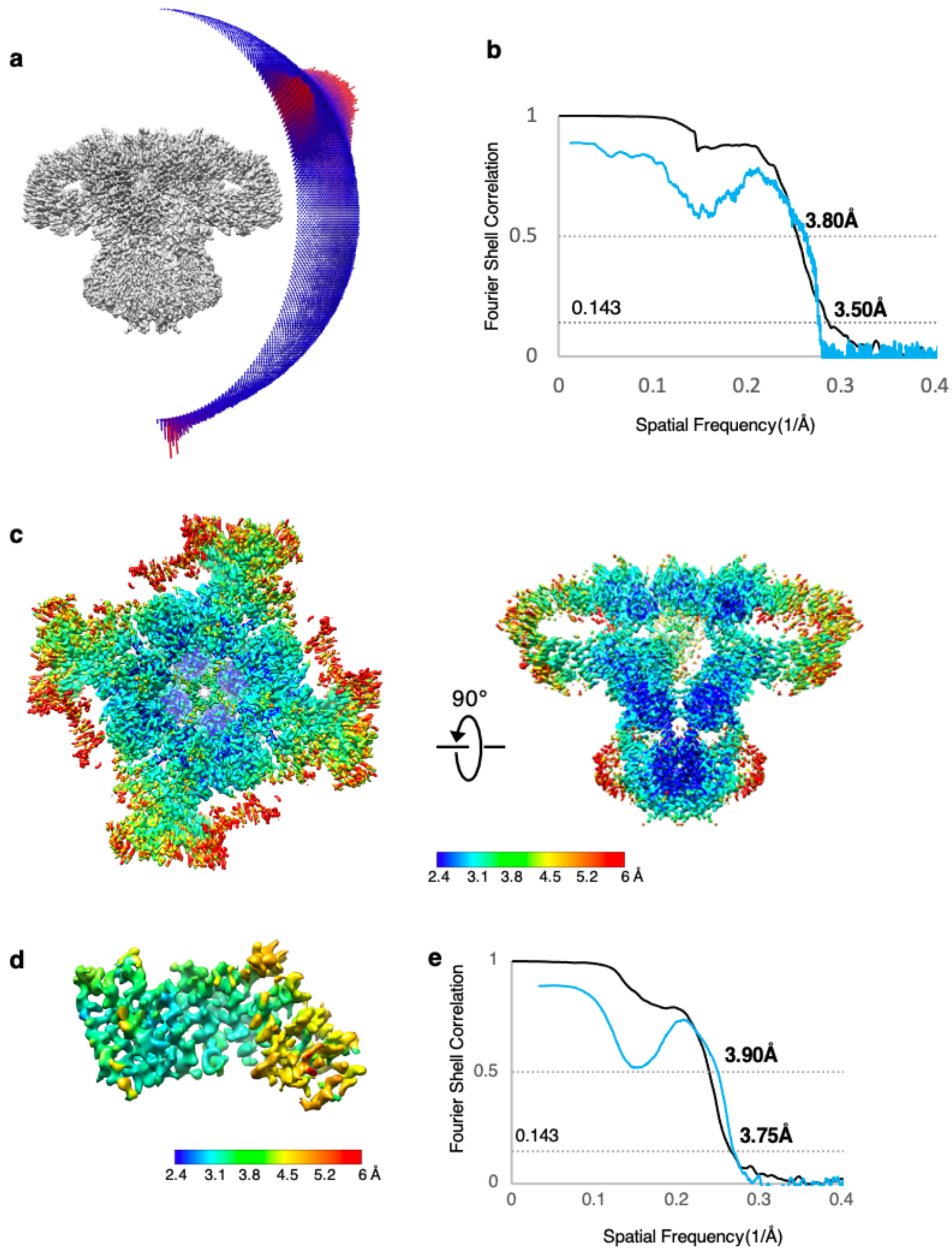


**Supplementary Figure 4. Characteristics of cryo-EM reconstructions for CIA-IP$_3$R1**. **a,** Euler angle distribution of particles contributing to the final reconstruction with larger red cylinders representing orientations comprising more particles. **b,** Resolution estimation of the cryo-EM density maps. Black – gold-standard Fourier shell correlation (FSC) curves, showing the overall nominal resolutions of 3.5 Å at 0.143 criterion; blue - Map-model FSC plot calculated by Phenix. **c,** Cryo-EM map colored based on local resolution. **d,** Focused 3D refined ARM2 map colored based on local resolution. **e**, Resolution estimation of the focused 3D refined ARM2 domain density map. Black – gold-standard Fourier shell correlation (FSC) curves, showing the overall nominal resolutions of 3.7 Å at 0.143 criterion; blue - Map-model FSC plot calculated by Phenix.

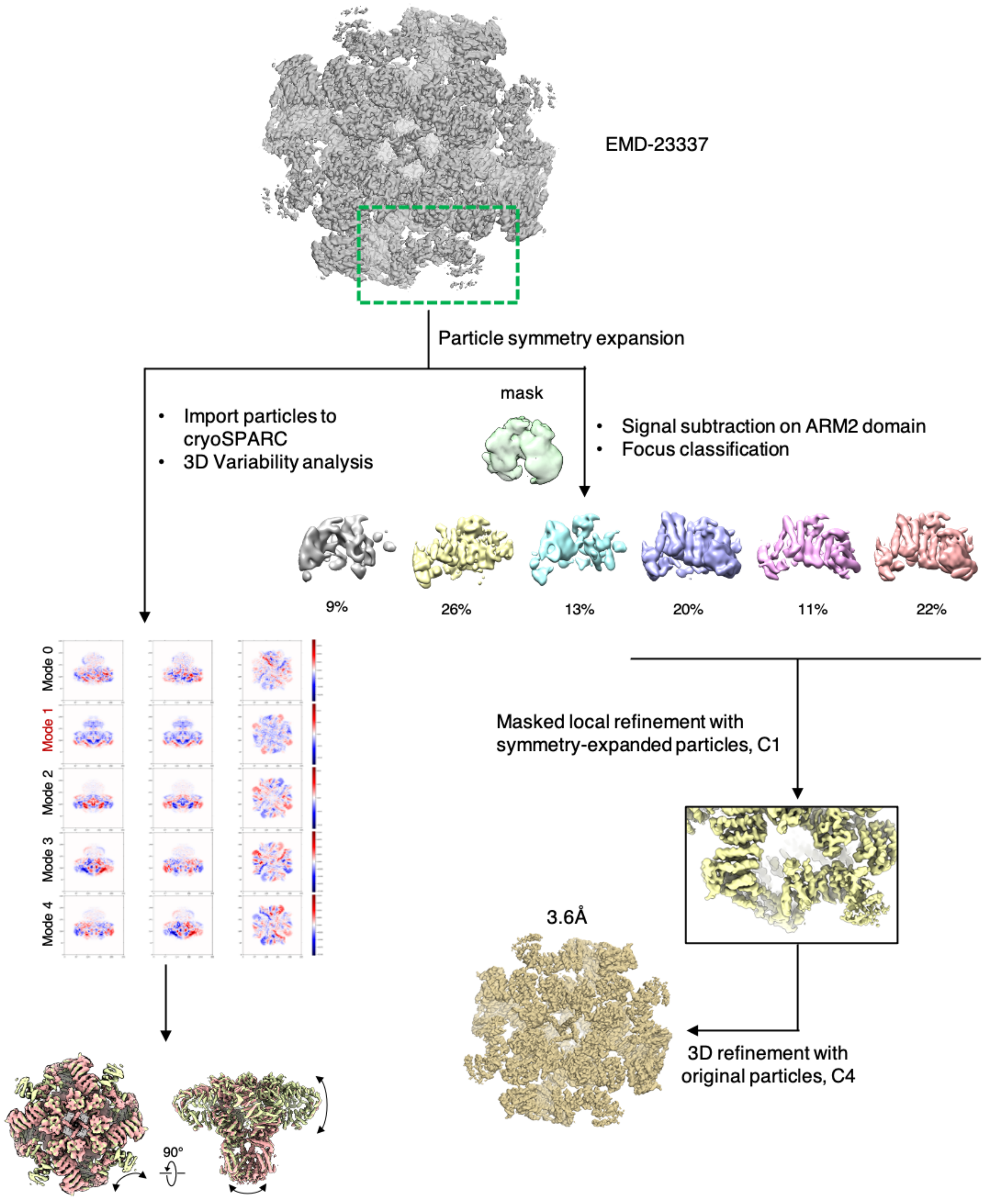


**Supplementary Figure 5.** 3D focused classification and local refinement on ARM2 domain for Apo-IP$_3$R1 in nanodisc used in Baker *et al.*[27]

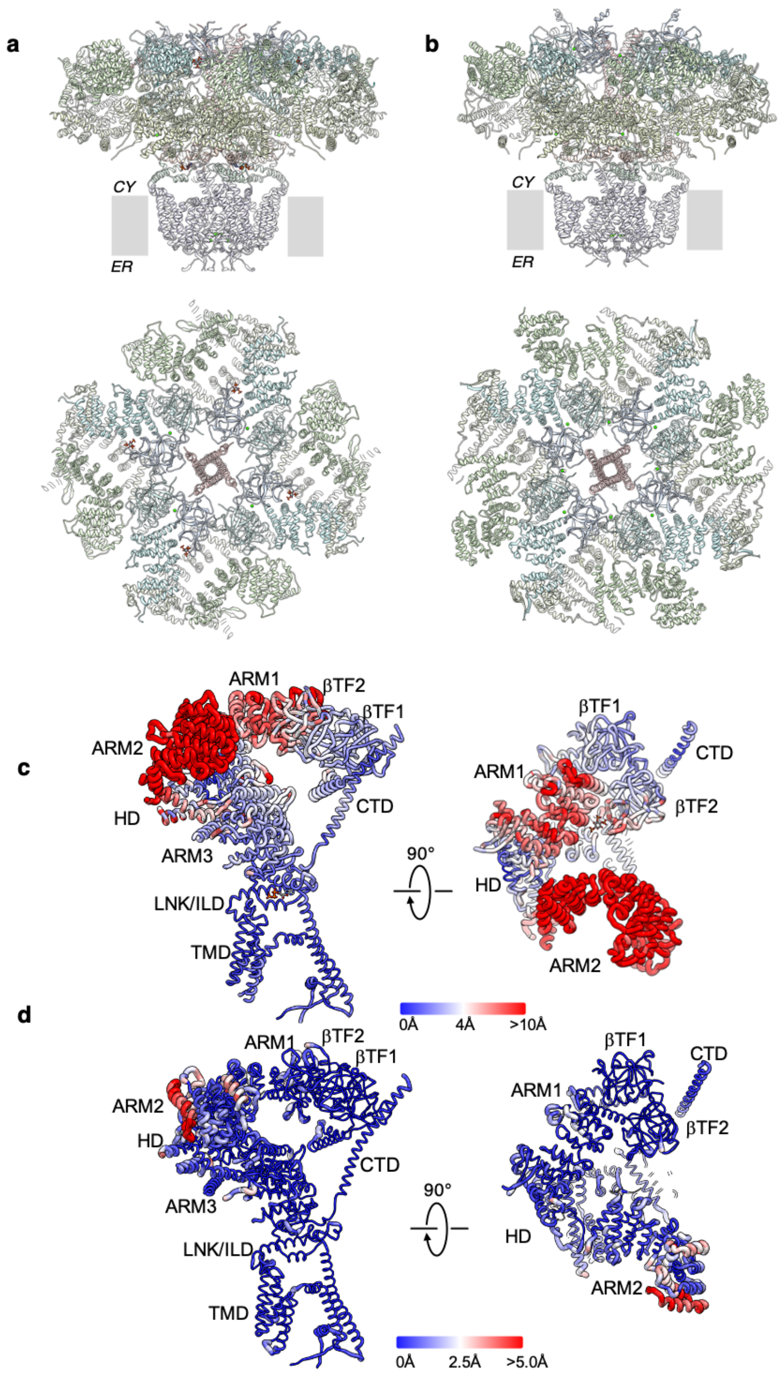


**Supplementary Figure 6.** Atomic models for the full-length tetrameric assembly of CIA-$IP_3R1$ (**a**) and Ca-$IP_3R1$ (**b**) based on the respective cryo-EM density maps. Top panels show side views along the membrane plane. Transmembrane boundaries are indicated in gray. Bottom panels show top-views from the cytosol along the four-fold axis. **b**, $C\alpha$ RMS deviations were calculated between CIA-$IP_3R1$ and Ca-$IP_3R1$ (**c**) and Ca-$IP_3R1$ and apo-$IP_3R1$ (PDB ID: 7LHE) (**d**) resulting in 11 Å and 2 Å RMSD, respectively. Per-residue deviations are mapped onto one subunit and color-coded based on their deviation lowest RMSD (blue/thinnest) to highest RMSD (red/thickest) is reflected in the ribbon color/thickness.

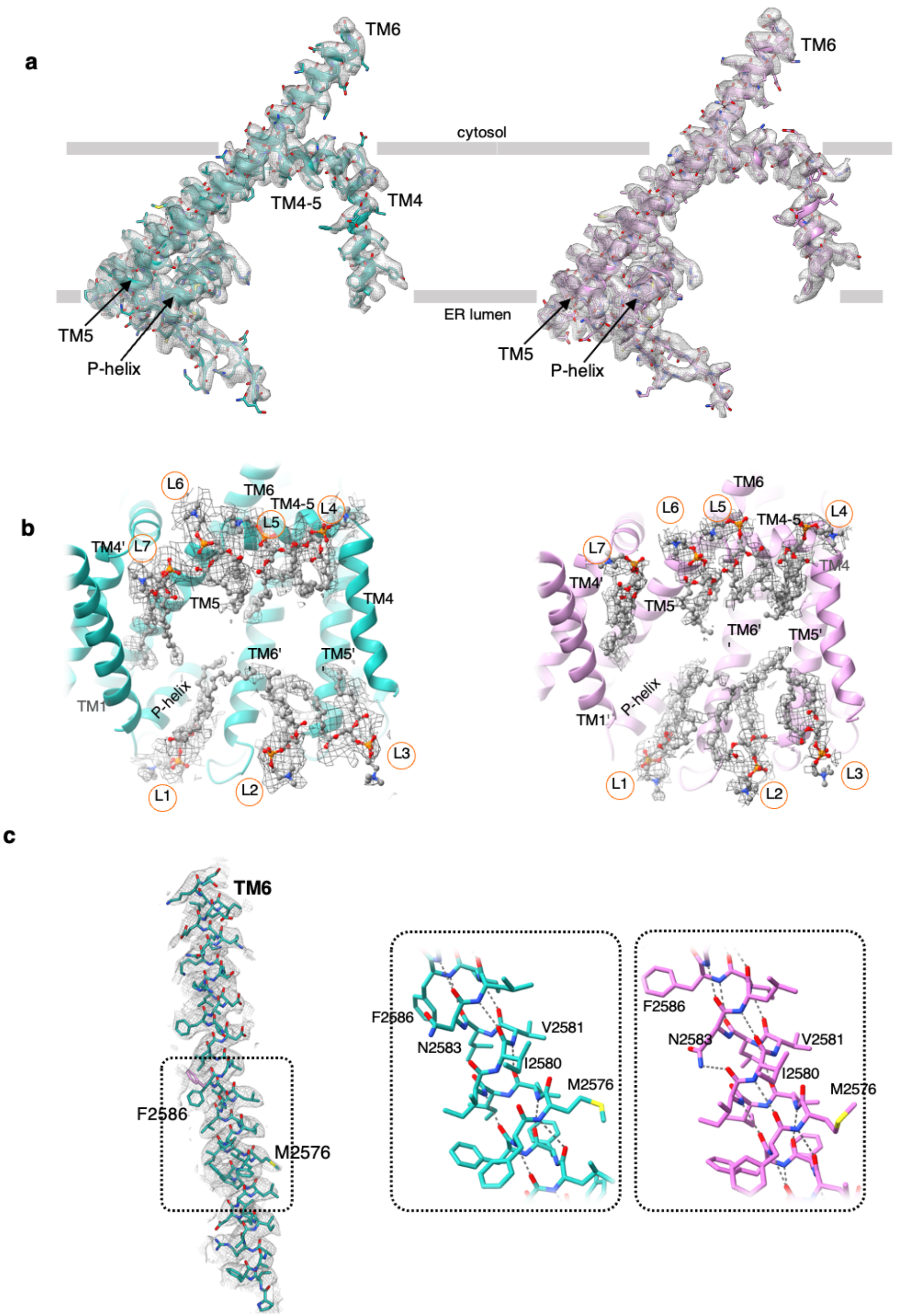


**Supplementary Figure 7. Structural features in the IP$_3$R1 transmembrane domains. a,** Densities (gray) for TMDs TM4 through TM6 from one subunit are overlaid with their atomic models CIA-IP$_3$R1 (blue-green), Ca-IP$_3$R1 (pink) and membrane boundaries indicated. **b,** Non-protein densities (gray mesh) within the transmembrane domains corresponding to lipids were identified in CIA-IP$_3$R1 (left) and Ca-IP$_3$R1 (right) are consistent with those described in Baker et al.[27]. The model of the TMDs is depicted as a ribbon and viewed parallel to the membrane plane. Lipids are represented as ball-and-stick models and labeled L1–L7. **c,** TM6 helix in CIA-IP$_3$R1 (blue-green) is overlaid with cryo-EM densities, colored gray (oxygen atoms - red; nitrogen atoms - blue; sulfur atoms - yellow); boxed region indicates the location for the $\pi$-helix in TM6. Zoomed-in views of the $\pi$-helix in CIA-IP$_3$R1 (blue-green) and Ca-IP$_3$R1 (pink) are shown in inserts.

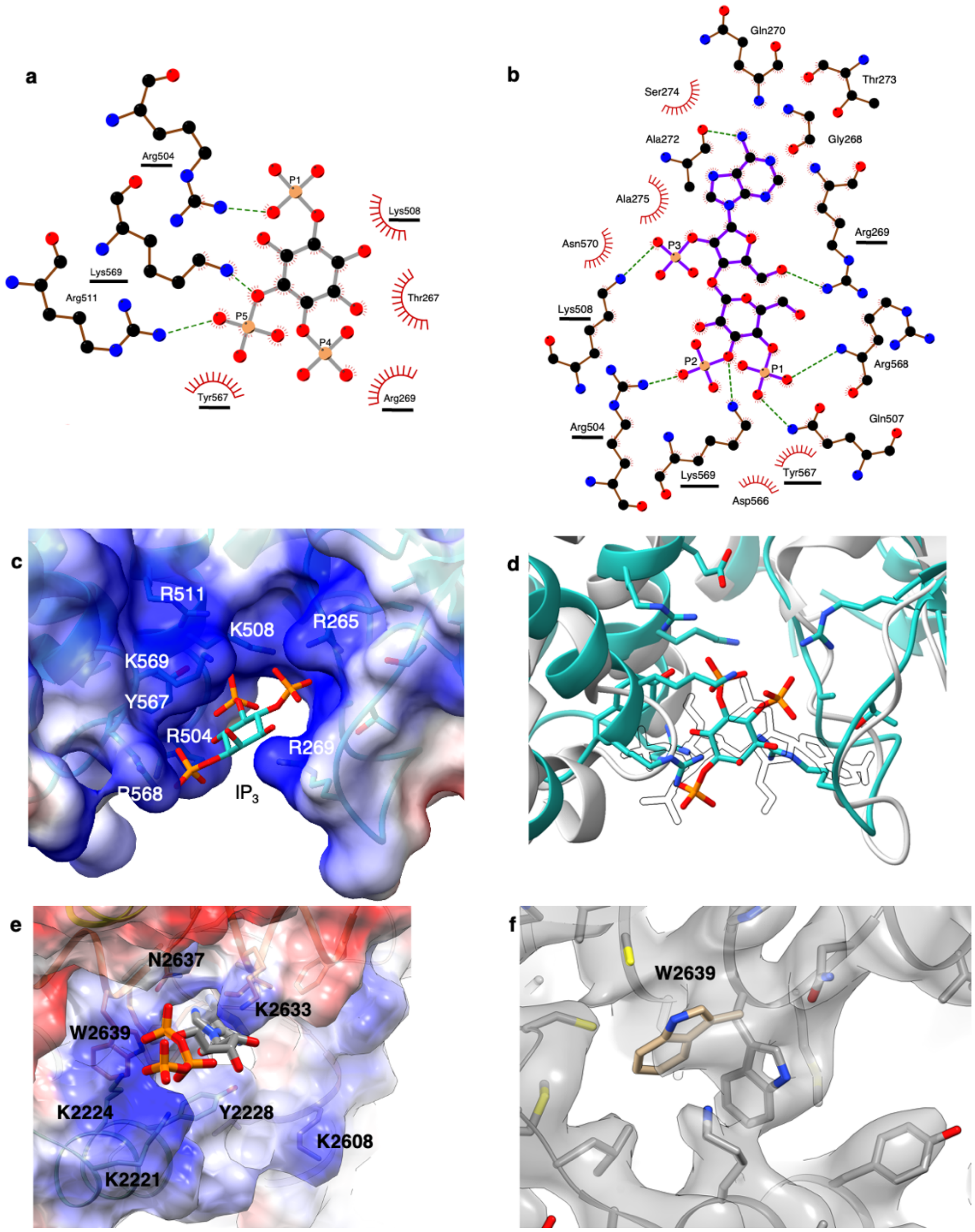


**Supplementary Figure 8. Properties within the $IP_3$ and ATP ligand binding pockets. a,** Schematic plot of the $IP_3$ molecule interacting with surrounding residues within the in CIA-$IP_3$R1 structure. **b,** Schematic plot of the Adenophostin A molecule interacting with surrounding residues in the ADA-$IP_3$R1 (PDB ID: 6MU1) structure. $IP_3$R1 residues that are consistent in coordinating ADA and $IP_3$ are underlined. Arcs represent hydrophobic interactions, green dashed lines are H-bonds as calculated in LigPlot+[93]. **c**, Coulombic electrostatic potential calculated and displayed on the surface representation of the $IP_3$ binding pocket in CIA-$IP_3$R1 with the pocket lining residues labeled. Blue indicates positive, red indicates positive. **d,** Structural alignment and overlay of the $IP_3$ binding pocket in CIA-$IP_3$R1 (blue-green) and ADA-$IP_3$R1 (PDB ID: 6MU1, white). **e**, Coulombic electrostatic potential calculated and displayed on the surface representation of the ATP binding pocket in CIA-$IP_3$R1 with the pocket lining residues labeled. **f**, Zoomed-in view of the ATP binding pocket in apo-$IP_3$R1 (EMDB: 23337) overlaid with its molecular model (PBD ID: 7LHE) and focused on the side chain conformers for W2639.

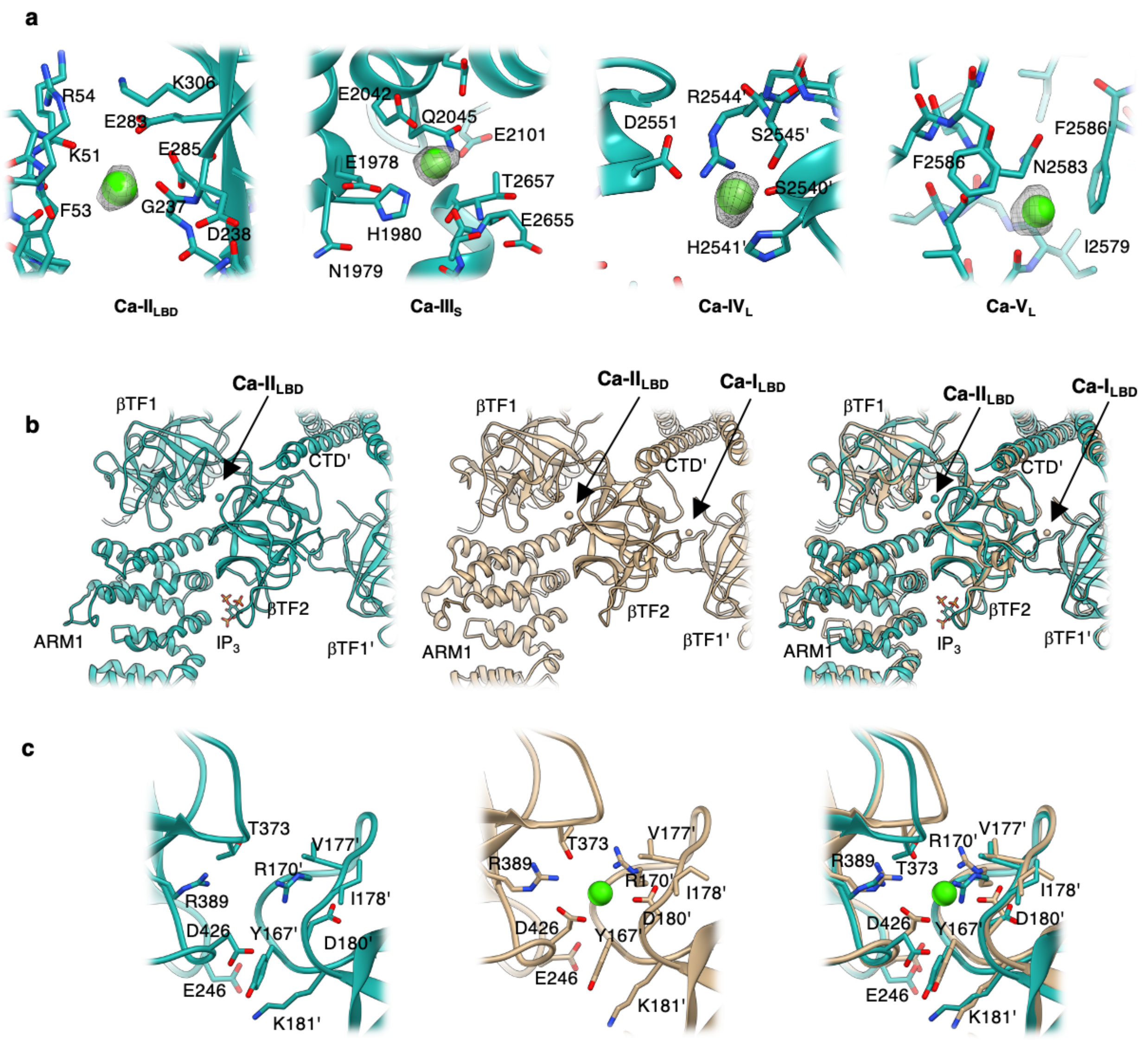


**Supplementary Figure 9. $Ca^{2+}$ binding sites identified in CIA-IP$_3$R1. a,** $Ca^{2+}$ binding sites localized in the ligand binding domains (Ca-II$_{LBD}$), the $Ca^{2+}$ sensor/ARM3 domain (Ca-III$_S$), and luminal vestibule of the TMD (Ca-IV$_L$ and Ca-V$_L$). $Ca^{2+}$ ions are shown as green spheres and overlaid with corresponding densities displayed at 2-5 $\sigma$ cutoff values. Residues within 5 Å of the $Ca^{2+}$ ions are displayed in a stick representation and labeled. **b**, Comparison of $Ca^{2+}$-binding sites identified in ligand binding domains in CIA-IP$_3$R1 (green) and Ca-IP$_3$R1 (tan). **c**, Zoom-in views of Ca-I$_{LBD}$ binding sites in CIA-IP$_3$R1 (green) and Ca-IP$_3$R1 (tan).
